# Repression precedes the stepwise evolution of a highly specific gene expression pattern

**DOI:** 10.1101/2020.11.11.378737

**Authors:** Jian Pu, Zinan Wang, Haosu Cong, Jacqueline S.R. Chin, Jessa Justen, Joanne Y. Yew, Henry Chung

## Abstract

Well-controlled gene expression is critical for the proper development and function of many traits. Highly-specific temporal and spatial expression patterns are often due to the overlapping activities of activator and repressor sequences that form *cis*-regulatory elements called enhancers. While many studies have shown that evolutionary changes in enhancers can result in novel traits, few studies illuminate how enhancers originate, how activator and repressor sequences interact during enhancer evolution, and the order in which they evolve. Here, we traced the evolutionary origin of a recently evolved enhancer that drives the expression of the fatty acyl-CoA elongase, *bond*, specifically in the semicircular wall epithelium (swe) of the *Drosophila* male ejaculatory bulb (EB). We show that this enhancer consists of two activator regions that drive *bond* expression in the entire EB and a repressor region that restricts expression specifically to the EB swe. Interestingly, the repressor region preceded the evolution of the two activator regions. The evolution of the first activator region, consisting of two putative *Abdominal-B* sites, did not drive expression in the EB due to the action of the repressor region. Expression of *bond* in the EB swe requires the evolution of the second activator region, which does not drive expression on its own, but synergizes with the first activator region and the repressor region to produce a highly-specific spatial expression pattern. Our results show that the origin and evolution of a novel enhancer require multiple steps and the evolution of repressor sequences can precede the evolution of activator sequences.

## Introduction

Highly-specific gene expression patterns are central to the development and evolution of multicellular organisms. These expression patterns are controlled by the action of modular *cis*-regulatory elements called enhancers (1). Enhancers are generally made of positive regulatory elements, such as activator sequences that recruit transcriptional activators to drive expression of a gene, and negative regulatory elements, such as repressor sequences that recruit transcriptional repressors to inhibit gene expression (1–3). The overlapping and combinatorial actions of activators and repressors within an enhancer control highly-specific gene expression in time and space (4).

A well-studied example of this logic is the even-skipped (*eve*) stripe 2 enhancer, where the combinatorial actions of two activators and three repressors define a single stripe of *eve* expression in the developing insect embryo (1,5–7). In the past few decades, numerous studies have demonstrated that changes in enhancer activities can lead to novel expression patterns and evolutionary innovations in morphology and physiology (8–10). These evolutionary changes in the enhancers include the gain (11) and loss (12, 13) of activator sequences as well as the gain (14) and loss (15) of repressor sequences. While these studies clearly show that evolutionary changes in either activator sequences or repressor sequences can lead to phenotypic changes, fundamental questions remain regarding how these activator and repressor sequences interact during the origins of these highly-specific enhancers.

In one of the first studies of *cis*-regulatory changes and evolutionary novelty, the gain of a novel enhancer that underlies a highly-specific wing spot in males of *Drosophila biarmipes*, is hypothesized to be due to recruitment of at least one activator and one repressor (16–18). How do enhancers that rely on the combinatorial effects of both activator and repressor sequences evolve? Did activator sequences evolve first, or did repressor sequences evolve first? A reasonable hypothesis would be that activation sequences evolved first, driving a broad expression pattern prior to the appearance of repressor sequences that restricts expression of the gene. However, a caveat of this prediction is that for many pleiotropic genes, gains in broad expression patterns may lead to negative fitness effects in the organism, due to possible misexpression of this gene in key tissues. Many studies in model organisms such as *Drosophila*, mice, and zebrafish have shown that misexpression of key genes in the wrong tissues can lead to negative phenotypic effects on the organism (19–22). In humans, the mis-regulation of genes underlies many disease conditions (23). Therefore, these negative fitness effects may lead to new broad expression patterns being selected against before the gain of repressor sequences.

An alternate hypothesis is that repressor sequences precede the evolution of activator sequences, leading to the gain of a specific expression pattern without having evolved broad expression first, thus mitigating the negative fitness effects of broad gene expression patterns. However, the hypothesis also poses a potential quandary: Can repressor sequences evolve before activator sequences if no gene expression is driven by repressors alone? Determining the evolutionary dynamics between activator and repressor sequences can lead to a better understanding of how enhancers originate and how highly-specific expression patterns in nature are generated during evolution.

In this study, we investigated the *cis*-regulatory evolution of the fatty acyl-CoA elongase gene *bond*, a pleiotropic gene involved in *Drosophila* spermatogenesis (24) as well as the biosynthesis of the male anti-aphrodisiac, CH503, a male-specific pheromone produced by several *Drosophila* species (25, 26). The production of CH503 requires the specific expression of *bond* in the male ejaculatory bulb (EB) (26). Using GFP reporter assays in transgenic *D. melanogaster*, we showed that *bond* is specifically expressed in the EB semicircular wall epithelium (swe). In addition, we show that *bond* expression is controlled by an enhancer consisting of two activator regions that drive *bond* expression in the entire EB and a repressor region that restricts expression specifically to the EB swe. Interestingly, this repressor region is present even in distantly related *Drosophila* species, *D. willistoni* and *D. virilis*, where *bond* is not expressed in the EB of either species. Further experiments show that the one of these species, *D. willistoni*, has evolved activator sequences that can drive reporter gene expression partially in the EB handle base. However, *bond* is not expressed in the EB of this species due to the action of the repressor region. Taken together, our experiments suggest that the evolution of the *bond* EB enhancer is stepwise, and the repressor sequences precedes the gain of activator sequences in the evolution of this highly specific enhancer.

## Results

### *cis*-regulatory evolution underlies the differential expression of *bond* in different *Drosophila* species

To investigate how the expression of *bond* in EB arose, we first sought to locate the enhancer driving this expression pattern in *D. melanogaster*. GFP reporter constructs using non-coding DNA sequences around the *bond* gene in *D. melanogaster* showed that the enhancer responsible for driving gene expression in the EB lies in the first intron of *bond* (**Fig. 1A, S1**). The EB is made of distinct parts including a bulbar cavity devoid of cells which holds the ejaculate (27)(**Fig. 1B**). GFP expression in the EB is restricted to a specific epithelial region which we named the semicircular wall epithelium (swe) (**Fig. 1B, C**), consistent with expression detected by *in situ* hybridization (**Fig. S2**) and previous observations (26). Homologous regions from six other *Drosophila* species driving GFP expression in *D. melanogaster* showed that the sequences from other *melanogaster* group species *D. simulans, D. yakuba, D. erecta*, and *D. ananassae* drove similar expression in the EB swe. However, the first intron of *bond* in distantly-related species, *D. willistoni* and *D. virilis*, did not drive expression in the EB at all (**Fig. 1C**). This result is consistent with previous expression studies of *bond* in some of these species (26). Our current experiments show that this difference in the expression of *bond* in the EB across *Drosophila* species is due to the evolution of *cis*-regulatory sequences present in the first intron of *bond*.

**Figure 1.**
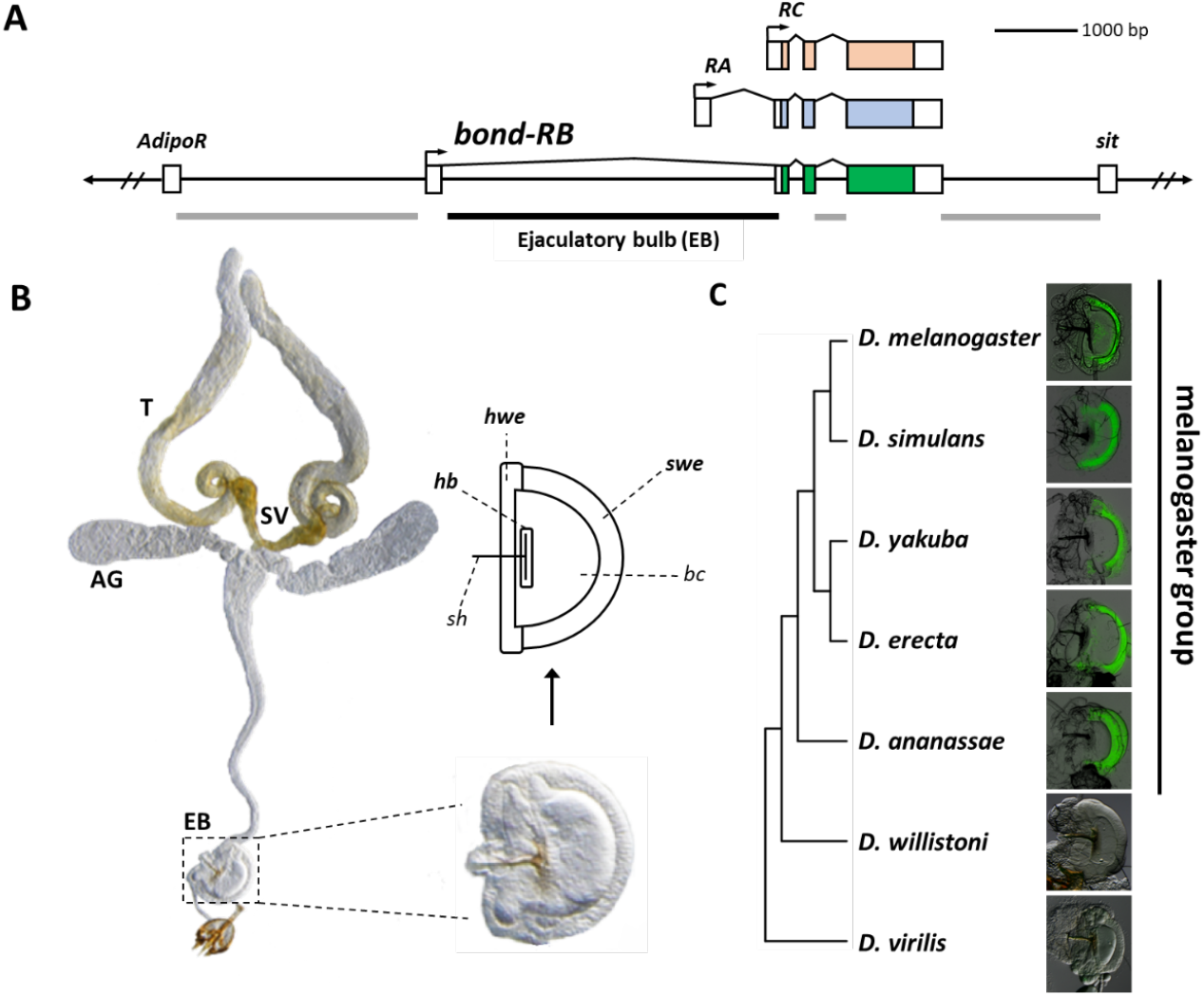
*Cis*-regulatory evolution underlies the ejaculatory bulb (EB) expression of *bond* in different *Drosophila* species. **(A)** The non-coding regions around the *D. melanogaster bond* locus was screened for enhancer activities that were able to drive GFP reporter protein expression in the EB. One fragment (black line), the first intron, was able to drive GFP expression in the EB. The other fragments (grey lines) did not drive EB expression. **(B)** Schematic of the male reproductive system in *D. melanogaster*. T = testes, AG = accessory gland, SV = seminal vesicle, EB = ejaculatory bulb. Parts of the EB: *hwe* = horn wall epithelium, *swe* = semicircular wall epithelium, *bc* = bulbar cavity, *hb* = handle base, *sh* = sclerite handle. **(C)** Intron 1 of *bond* from *D. melanogaster, D. simulans, D. yakuba, D. erecta*, and *D. ananassae* drove GFP expression in the semicircular wall epithelium of the EB. The homologous fragments from *D. willistoni* and *D. virilis* did not.

### A 285 bp enhancer, comprised of both activator and repressor sequences, recapitulates specific expression of *bond* in the semicircular wall epithelium (swe) of the *D. melanogaster* EB

Having confirmed that *cis*-regulatory differences underlie *bond* expression differences in the swe of the EB, we sought to identify the evolutionary changes in the *cis*-regulatory sequences that led to the differential expression patterns. Our first step was to determine the minimum enhancer region specific for *bond* expression in the EB swe. We performed a systematic dissection of the first intron of *D. melanogaster* to delimit smaller regions for the enhancer by creating smaller overlapping GFP constructs. We identified a 285bp region (construct *bc23*) that recapitulated the expression pattern driven by the full intron **(Fig. 2A, S1)**, We named this the EB semicircular wall epithelium (swe) enhancer. We next set out to identify sequences in this minimal enhancer that potentially underlie *bond* expression differences between species. We initially divided the 285bp enhancer fragment into overlapping constructs, *bc2* and *bc3. bc2* did not drive any GFP expression, but notably, *bc3* drove expression in the entire EB, not just the swe of the EB **(Fig. 2A, B)**. This result suggests that repressor sequences present in the *bc2* fragment can repress the expression in the horn wall epithelium (hwe) and handle base (hb) of the EB driven by the *bc3* fragment **(Fig. 1B)**. Dissection of *bc3* into two smaller overlapping constructs, *bc3i* and *bc3ii*, showed that while *bc3i* drove expression in the hb, *bc3ii* did not drive any GFP expression. This result suggests that *bc3ii* contains activator sequences necessary to drive expression in the whole EB in conjunction with *bc3i*, but cannot independently drive expression. Together, these data show that the *bond* EB swe enhancer is divided into at least one repressor region (Rep) and two activator regions (Ac1 and Ac2) **(Fig. 2C)**. Additional experiments showed that this repressor is modular and can repress the EB enhancer of another gene in a distance-dependent manner **(Fig S3)**.

**Fig. 2.**
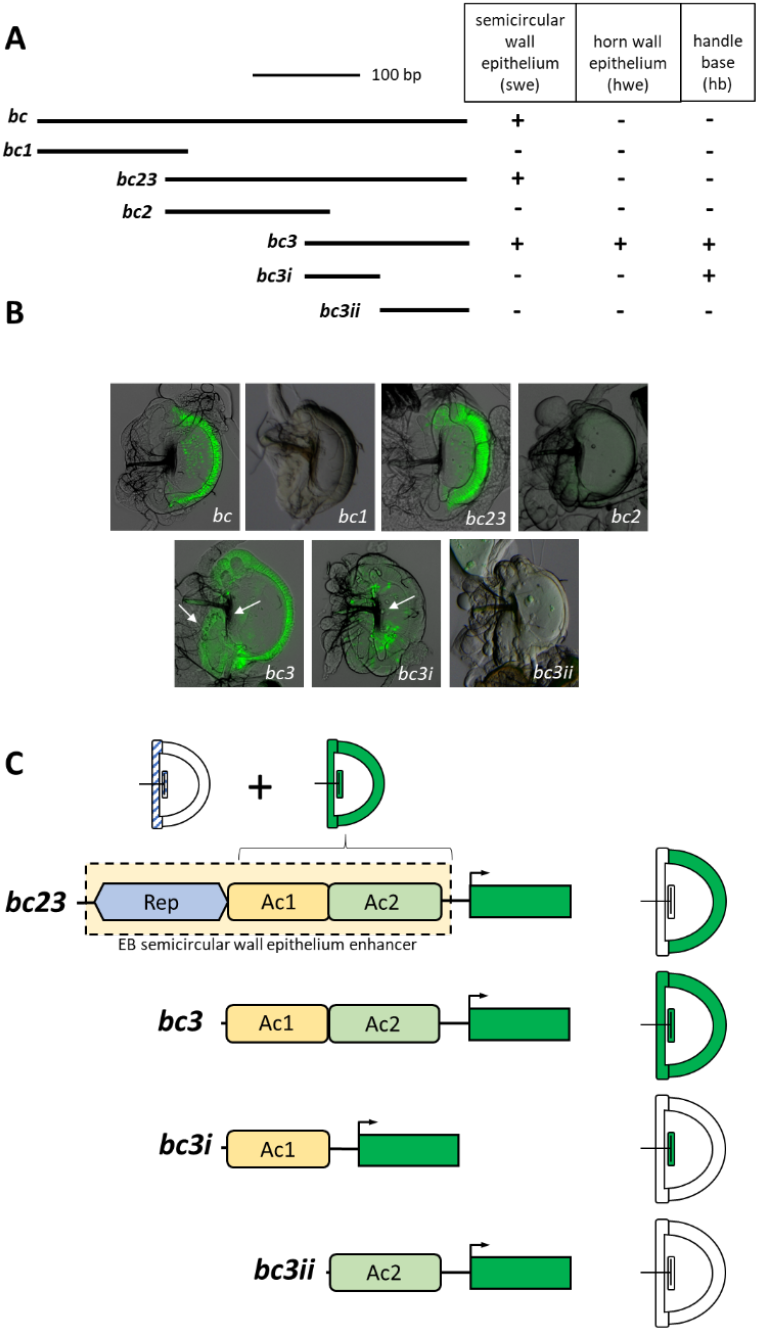
The combination of repressor and activator sequences in the *D. melanogaster bond* EB enhancer drives specific expression in the semicircular wall epithelium. (A) Schematic of the overlapping GFP constructs of the EB enhancer. The bc23 fragment is the minimum region that can recapitulate the EB expression of *bond* in the semicircular wall epithelium. The bc3 fragments shows the ectopic GFP expression in horn wall epithelium (hwe) and handle base (hb). (+) indicates the presence of expression, and (-) indicates the absence of detectable expression. (B) GFP reporter protein expression in the EB corresponding to the different overlapping constructs. Arrows indicate the expression in hwe and hb. (C) Model for the *D. melanogaster bond* EB enhancer. There are three modular regions in the enhancer. The Ac1 region contains activator sequences that drive expression in the hb; the Ac2 region contains activator sequences to drive expression in the entire EB (in hwe, hb and swe) in combination with the Ac1 region, but not drive GFP expression on its own; the repressor region Rep represses the activity of the Ac1 and Ac2 regions in hb and hwe of EB and restricts their activity to the swe of the EB.

### Activator sequences for EB expression are present in *D. willistoni*, even though *bond* is not expressed in the EB of this species

To trace the evolutionary origins of this enhancer in *Drosophila*, we examined the activity of the homologous sequences from three other species, *D. ananassae, D. willistoni*, and *D. virilis*, based on their *bond* expression in the EB and their phylogenetic relationships **(Fig. 1C)**. Our results indicate that the *bc* fragments from these three species are able to recapitulate the EB expression of *bond* (in the case of *D. willistoni* and *D. virilis*, no expression). Our *a priori* expectation is that creating smaller fragments of the 285bp enhancer in *D. melanogaster* and *D. ananassae* would allow us to narrow down the region involved in enhancer evolution, and we expected that there would be no GFP expression driven by the smaller fragments in *D. willistoni* and *D. virilis*. While we did not detect GFP expression in any of the *D. virilis* constructs, to our surprise, the *bc3i* fragment of *D. willistoni* was able to drive GFP expression in the hb of the EB, similar to homologous fragments in *D. melanogaster* and *D. ananassae* **(Fig. 3A)**. This result suggests that there are activator sequences in *D. willistoni* that can drive partial GFP expression in the EB. Our observation that the *D. willistoni bc3i* fragment can drive GFP expression in the hb of the EB but not the full *D. willistoni bc* fragment suggests that repressor sequences similar to the repressor region present in *D. melanogaster* may also be present in *D. willistoni* **(Fig. 3A)**.

**Fig. 3.**
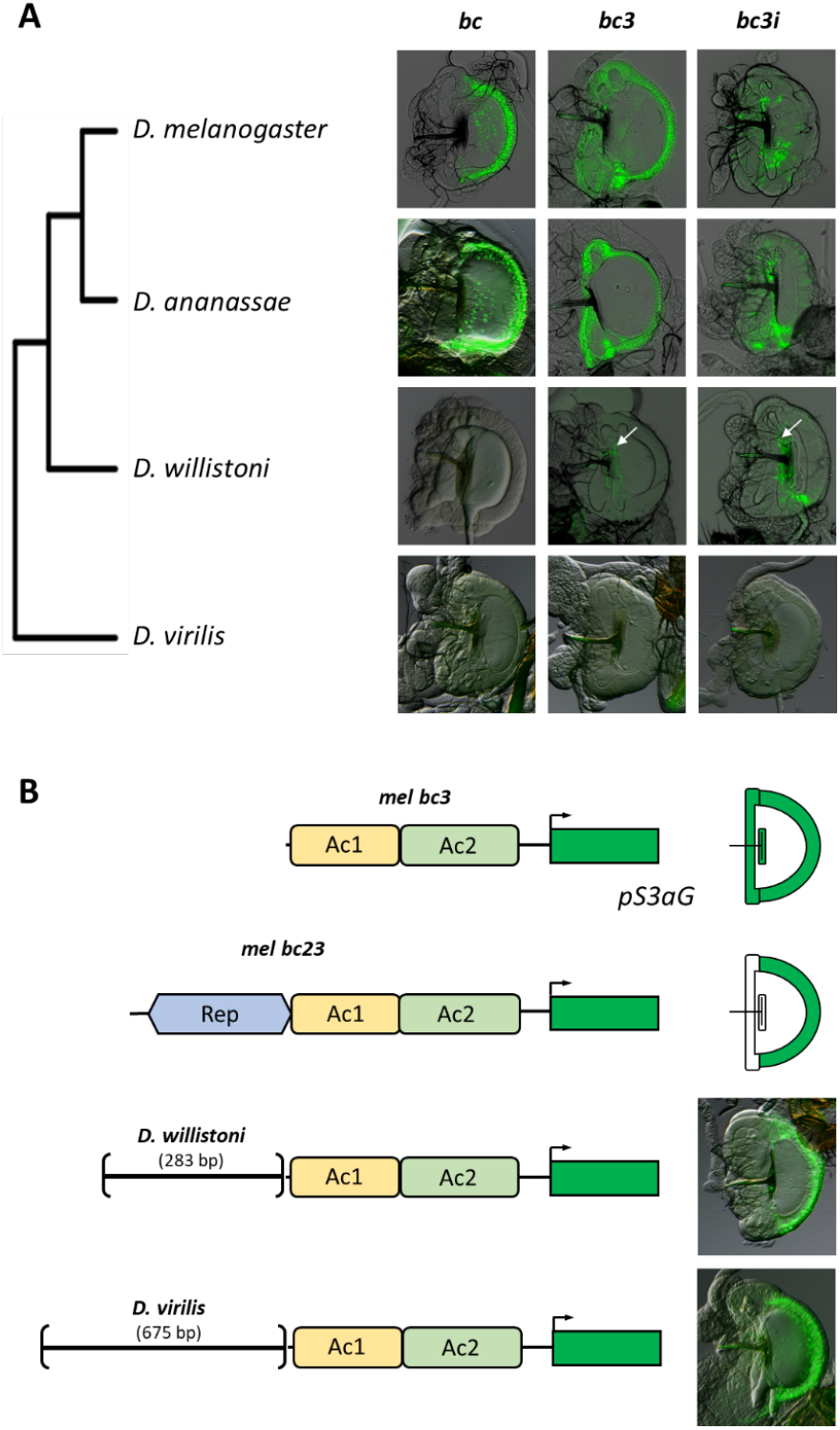
Repressor sequences that can repress *bond* expression in the EB hwe and hb are present in all species tested, including species that do not express *bond* in the EB. (A) The larger *bc* construct recapitulates the native expression of *bond* in the EB in *D. melanogaster* and *D. ananassae* and showed no EB GFP expression with *D. willistoni* and *D. virilis* homologous constructs, similar to their native expression. However, the smaller *D. willistoni bc3* and *bc3i* constructs could drive GFP expression in the handle base (white arrows), similar to the *bc3i* construct of *D. melanogaster*, suggesting the presence of repressor sequences in the larger bc construct of this species. (B) Homologous sequences to the *D. melanogaster* Rep region from *D. willistoni* and *D. virilis* can repress GFP expression in the hwe and hb driven by the D. *melanogaster bc3* construct, suggesting that these fragments contain repressor sequences similar to the *D. melanogaster* Rep region.

### Repressor sequences are present in species that do not express *bond* in the EB

To confirm our observation that repressor sequences may be present in *D. willistoni*, we created GFP reporter constructs that fused the region from *D. willistoni* homologous to the repressor region in *D. melanogaster* with the *D. melanogaster bc3* construct that drives expression in the entire EB. If the D. *willistoni* sequence functions as a repressor, it should spatially repress expression of the *D. melanogaster bc3* fragment and restrict expression to the swe of the EB in *D. melanogaster*. Our results confirm this prediction: The *D. willistoni* fragment effectively repressed *bc3* driven GFP expression in the hwe and the hb, and restricted expression to the swe, thus confirming that repressor sequences are present in *D. willistoni* **(Fig. 3B)**. We tested the homologous regions from another species, *D. virilis*, where *bond* is not expressed in the EB. Intriguingly, these homologous regions can also repress expression driven by the *D. melanogaster bc3* construct, restricting GFP expression in the EB swe. Taken together, these observations suggest that repressor sequences are present in *D. willistoni* and *D. virilis* and precede the evolution of the complete minimal EB swe enhancer. To narrow down the sequences involved in repression, we made smaller constructs of the *D. melanogaster* 141bp Rep region and assayed for their abilities to repress GFP expression driven by the *bc3* construct (Ac1 and Ac2 regions) (**Fig. S4**). These constructs, coupled with additional site-directed mutagenesis experiments, narrowed the repressor sequence to an 11-bp sequence (5’-aaattaattta-3’), that is able to recapitulate the repressor activity of the *D. melanogaster* Rep region (**Fig. S4**). A search on Flybase (28) showed that there are 638 hits to this sequence in the *D. melanogaster* genome, implying the non-uniqueness and pervasiveness of this sequence throughout the genome.

### Evolution of putative *Abd-B* activator binding sites is involved in the stepwise evolution of *bond* expression in the EB

We have established that repressor sequences are present in the *bond* enhancer homologous sequences of two species that do not express *bond* in the EB. One of these species, *D. willistoni*, has sequences homologous to the activator region (Ac1) in *D. melanogaster* that could drive partial expression in the EB when isolated from the repressor, similar to that of *D. melanogaster* and *D. ananassae*. (**Fig. 3A**). This suggests that the gain of *bond* expression in the EB swe could be a stepwise evolutionary process. To investigate how the EB expression of *bond* evolved, we sought to identify potential transcriptional activators in this enhancer that can drive expression in the EB. We performed high throughput transcriptomic sequencing of EBs from 8-day old male *D. melanogaster* (**Table S1**) and identified putative transcription factors (TFs). Next, we carried out a reverse genetic screen by using an EB specific GAL4 driver (EB-GAL4) to drive *UAS::RNAi* constructs of the most highly expressed TFs (**Table S2**) in order to ascertain which of the TFs can affect *bc3*-driven GFP expression **(Fig. 4A, B)**. Out of the 100 *UAS::RNAi* constructs tested, one line, driving RNAi knockdown of the homeobox gene, *abdominal-B* (*Abd-B*), was able to disrupt *bc3*-driven GFP expression in the EB, suggesting that *Abd-B* may function as a transcriptional activator for the expression of *bond* in the EB **(Fig. 4B)**.

**Fig. 4.**
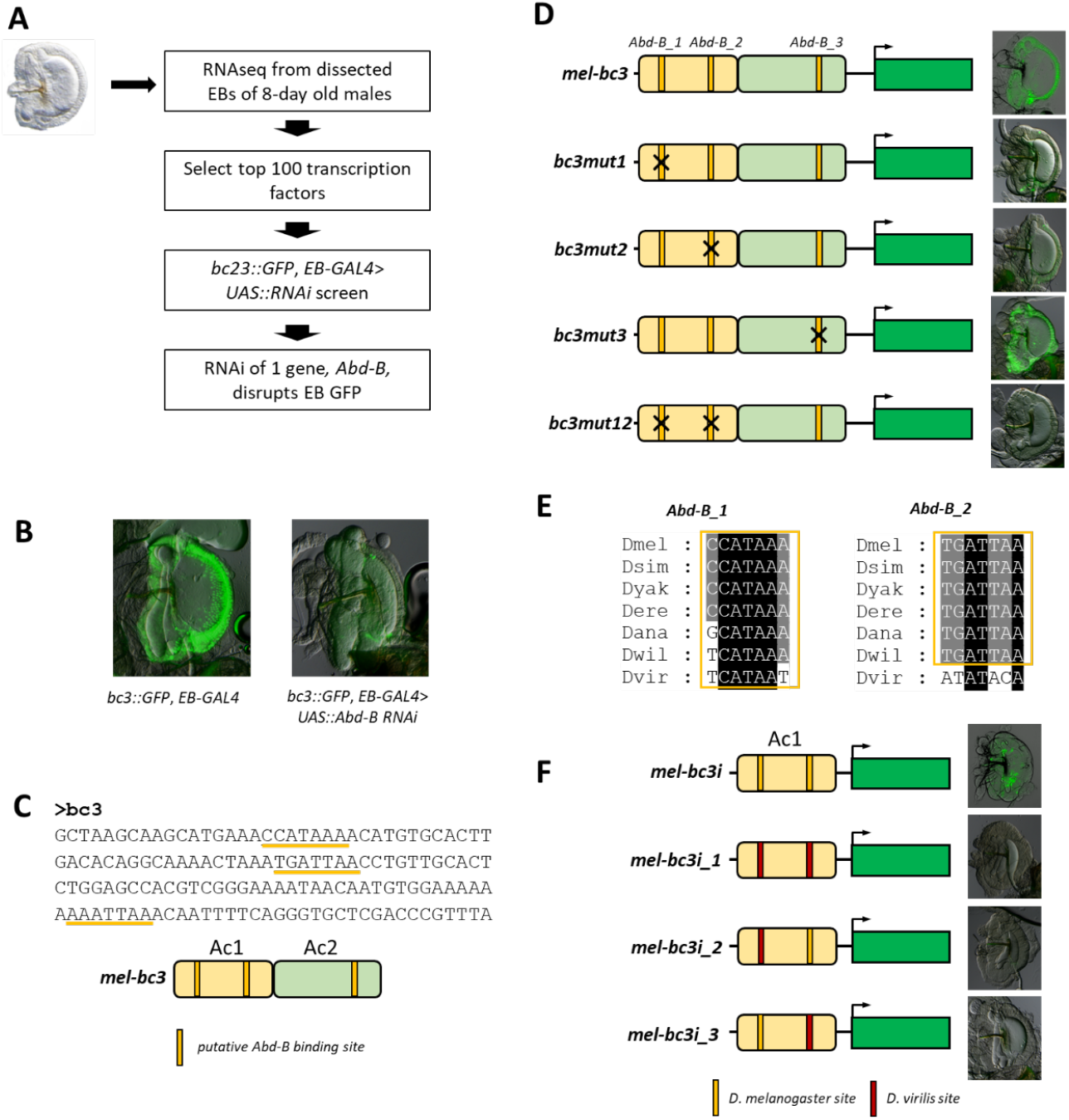
The evolution of two putative *Abd-B* binding sites is necessary for the expression of *bond* in the EB. (A) An EB RNAi screen identifies Abd-B as a possible regulator of *bond* in the EB. (B) RNAi knockdown of *bond* in the EB led to significant decrease in GFP expression driven by the *D. melanogaster bc3* construct. (C) JASPAR analysis identified three putative *Abd-B* binding sites in the *D. melanogaster bc3* fragment. (D) Site-directed mutagenesis of these putative *Abd-B* binding sites individually shows that the *Abd-B_1* and the *Abd-B_2* sites are necessary for GFP expression driven by the *bc3* construct, but not the *Abd-B_3* site. (E) Evolutionary analysis of the *Abd-B_1* and the *Abd-B_2* sites shows that all species have putative *Abd-B* binding sequences at *Abd-B_1*, and most species have putative *Abd-B* binding sequences at *Abd-B_2* except *D. virilis*. (F) Swapping in the *D. virilis* sequence at both sites either individually or in combination led to the loss of GFP expression in the hb of the EB driven by the *D. melanogaster bc3i* construct.

Bioinformatic analysis using JASPAR (29) predicted three putative *Abd-B* binding sites **(Fig. 4C)**. To determine whether the putative *Abd-B* binding sites function *in vivo*, we systematically mutated each site in the *bc3* GFP construct **(Fig. 4D)**. Our experiments show that site-directed mutagenesis of the first two putative *Abd-B* binding sites, *Abd-B_1* and *Abd-B_2*, reduced GFP expression drastically when mutated individually and completely abolished GFP expression when mutated in combination. In contrast, mutation of the third putative *Abd-B* binding site, *Abd-B_3*, did not affect GFP expression **(Fig. 4D)**. Our results suggest that *Abd-B_1* and *Abd-B_2* are both necessary to drive *bond* expression in the EB of *D. melanogaster*. As *Abd-B_1* and *Abd-B_2* are both in the Ac1 region (bc3i construct), we expect that these two sites will be conserved in all *melanogaster* group species tested and *D. willistoni*, but not *D. virilis*. To trace the evolution of *Abd-B_1* and *Abd-B_2* in these species, we performed an alignment of the *Ac1* region of these species **(Fig. S5)**. We used JASPAR to predict potential *Abd-B* binding sites in all these species and found that all species have putative *Abd-B* binding sequences at *Abd-B_1*, and most species have putative *Abd-B* binding sequences at *Abd-B_2* except *D. virilis* **(Fig. 4E)**. These predictions are consistent with the activity of homologous *bc3i* fragments (Ac1 region) from the four species, in which *D. melanogaster, D. ananassae*, and *D. willistoni*sequences drove GFP reporter expression in the hb of the EB, but *D. virilis* did not **(Fig. 3A)**. To determine if the predicted difference at the *Abd-B* binding sites could underlie differences in expression driven by the *D. melanogaster* and *D. virilis bc3i* constructs, we swapped in the corresponding *D. virilis* sequences at the *D. melanogaster Abd-B_1* and *Abd-B_2* sites individually in the *D. melanogaster bc3i* construct. Our results show that swapping in the *D. virilis* sequence at both sites either individually or together led to the loss of GFP expression in the hb of the EB **(Fig. 4F)**. Taken together, the findings suggest that evolution in both *Abd-B_1* and *Abd-B_2* sites between these species could contribute towards the evolution of *bond* expression in the EB.

## Discussion

The generation of highly-specific gene expression patterns is often due to the combinatorial actions of activators and repressors that form a modular enhancer driving these expression patterns (1, 4). However, the evolutionary history that leads to the origin of these enhancers is not always clear. In this paper, we have identified and characterized an enhancer for the elongase gene *bond* that drives precise expression of *bond* in the male EB semicircular wall epithelium (swe) of *D. melanogaster* and other species of the *melanogaster* group **(Fig. 1C)**. This specific expression of *bond* has been shown to be necessary for the production of the male anti-aphrodisiac pheromone, CH503 (26).

Our experiments show that the EB swe enhancer is made up of at least one repressor region (Rep) and two distinct activator region (Ac1 and Ac2), restricting *bond* expression to the swe of the EB in *D. melanogaster* **(Fig. 2C)**. We showed that a Rep region containing sequences capable of repressing gene expression in parts of the EB is present in all *Drosophila* species tested in our study, and precedes the evolution of the two distinct activator regions **(Fig. 5)**. During the divergence of the *Sophophora* and *Drosophila* subgenus, the Ac1 activator region, containing putative binding sites for the transcription factor *Abd-B*, evolved in the *Sophophora* subgenus. However, as seen in *D. willistoni*, Ac1 was unable to drive expression of *bond* in the EB due to the overlapping action of the repressor sequences. The Ac2 activator region that evolved in the ancestor of the *melanogaster* group was able to drive expression in the entire EB in conjunction with Ac1, but was not able to drive expression on its own **(Fig. 2C)**. Together, the combinatorial action of the Rep repressor region, and the Ac1 and Ac2 activator regions form a modular enhancer driving specific expression in the semicircular wall epithelium of the EB **(Fig. 5)**. Our model infers the stepwise gain of Rep, Ac1 and Ac2 in this order in the evolution of the EB swe enhancer. Based on the phylogenetic tree, four other alternate models are also plausible **(Fig. S6)**. One model suggests that the EB swe enhancer is ancestral to all these species and that *D. willistoni* and *D. virilis* underwent independent losses of the activator sequences **(Fig. S6A)**. However, this is unlikely as this model would invoke parallel losses of Ac2 in these two species and another loss of Ac1 in *D. virilis*, although this possibility cannot be excluded. The other three models also infer that repression precedes the evolution of activator sequences in the evolution of this enhancer, but these models are also less likely due to the fact that they invoked multiple gain and losses **(Fig. S6B-D)**.

**Fig. 5.**
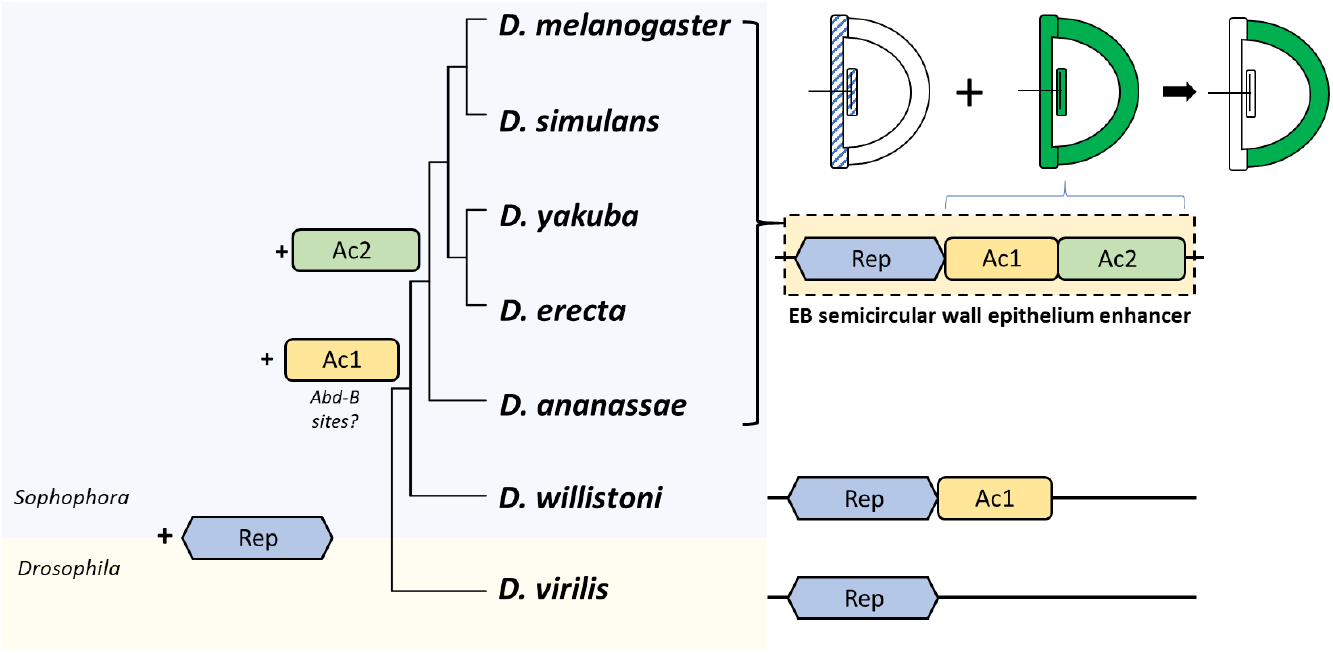
Repression precedes the stepwise evolution of a specific gene expression pattern. The EB semicircular wall epithelium (swe) enhancer is made up of two activator regions (Ac1 and Ac2) that drives expression in the whole EB (hb, hwe and swe), and a repressor region (Rep) that represses expression in the hb and hwe, resulting in the expression of *bond* in the EB swe. The Rep repressor region is present in all species and precedes the evolution of the activator sequences. The first activator region (Ac1) evolved in the Sophophora subgenus, possibly due to the gain of *Abd-B* binding sites. The Ac1 region is present in *D. willistoni* but there is no expression of *bond* in the EB of this species due to the presence of the Rep repressor region. The second activator region (Ac2) evolved in the *melanogaster* group and can drive expression in the whole EB in conjunction with Ac1, but due to presence of Rep, *bond* expression is restricted to the swe in these species.

### The evolution of modular enhancers

The evolution of modular enhancers driving novel expression patterns can occur from at least four different evolutionary mechanisms: transposition, promoter switching, co-option and *de novo* generation (8, 30). While the previous studies have provided evidence for the first three mechanisms, *de novo* evolution of enhancers has been difficult to study due to their rarity (8). Part of the challenge a *de novo* origin of an enhancer is due to the fact that these enhancers are usually composed of multiple bindingsites or sequences that bind different transcriptional activators or repressors. It is therefore more parsimonious to evolve a new enhancer by co-opting existing regulatory sequences (30). However, we argue that the *de novo* generation of an enhancer is possible. Previous genome-wide studies in *Drosophila* and humans suggest that a large fraction of these genomes are bound by transcription factors (31, 32). While there is a vigorous scientific discussion on whether these TF-bound sequences are functional (33, 34), the fact that TF-binding sites are short, degenerate, and dispersed widely and randomly across the genome suggests that a small number of mutations generating new TF binding sites around these existing TF-bound sequences would form a new enhancer based on the combinatorial effects of these sequences.

Our study shows that the gain of the Ac2 activator region in the *melanogaster* group led to the evolution of an EB semicircular wall epithelium enhancer. However, the Ac2 region is unable to drive expression on its own, relying on the combinatorial activity with the Rep repressor region and the Ac1 activator region to drive expression specifically in the EB swe. As we did not detect any other enhancer activities within 1kb of this region, we suggest that this could be a case of *de novo* generation of a novel enhancer, although we cannot rule out that co-option could also be a possibility (see discussion on the origin of the Rep region below).

### The role of repressor sequences in the evolution of highly specific expression patterns

Many previous studies have demonstrated the significance of repressor binding sites or sequences in modular enhancers to constrain and shape gene expression (14, 16, 35, 36). However, to our knowledge, there are no studies that investigate the order of whether activator sequences or repressor sequences evolved first.

The presence of repressor sequences (Rep) in *D. virilis* suggests that these sequences capable of repression are present before the divergence of the *Sophophora* and *Drosophila* subgenus (**Fig. 5**). What could be the putative function(s) of these sequences before being part of the *bond* EB swe enhancer? We propose three different hypotheses. The first hypothesis is that this repressor sequence is part of another enhancer in the *bond* intron, and is co-opted into the *bond* EB swe enhancer. However, our experiments in *D. melanogaster* showed that there are no other apparent enhancer activities within 1 kb of the Rep region aside from Ac1 and Ac2 (**Fig. S1**). In addition, our experiments suggest that the Rep region represses gene expression in a distance-dependent manner (**Fig. S3**), and is not able to repress gene expression that is more than 1 kb away. This could suggest that the repressor associated with the Rep region function as distance-dependent “short-range repressors” like knirps and Krüppel which can only repress gene expression around 100 bp away (1) rather than “long-range repressors” such as Hairy, which mediates repression of *cis*-regulatory sequences more than 1 kb away (37). A second hypothesis is the Rep region is evolutionarily conserved due to this region overlapping with the exon of an antisense non-coding RNA, *CR44062*, which resides on the opposite DNA strand to *bond* (**Fig. S7**). Although the function of *CR44062* is unknown, one possible scenario is that the Rep region is conserved due to potential functional constraint on the evolution of *CR44062*, i.e. sequence evolution of *CR44062* may have negative fitness effects. A third hypothesis is that binding sites for the transcriptional activators or repressors are usually very short (6-10 bp long) (38, 39). In our case, we narrowed down this repressor sequence to 11bp. Therefore, the likelihood of these short sequences randomly distributed across the genome without any apparent function is high. Bioinformatic analysis showed that this exact 11 bp sequence has 634 complete matches in *D. melanogaster* genome. This result suggests that some of the repressor sequences may be pervasive throughout the genome but do not produce any phenotypes until activator sequences that produce an overlapping expression pattern with the repressor sequences evolve. The phenomenon that repressor sequences preceding the gain of activator sequences in enhancer evolution could be very common during the evolution of highly-specific gene expression patterns. This may reflect a general evolutionary mechanism for enhancer origins.

## Materials and Methods

### Fly strains and transgenic constructs

All GFP reporter constructs were generated by PCR amplification of the genomic fragments from different *Drosophila* species and cloned into the GFP reporter vector *pS3aG* via the *AscI* and *SbfI* site (All primers listed in **Table S3**). All constructs were injected into the *Xout D. melanogaster* line and integrated into the genome using the *PhiC31* integrase system. The Canton-S strain was used as the wild-type *D. melanogaster* strain. *D. simulans, D. yakuba, D erecta, D. ananassae, D. willistoni*, and *D. virilis* were the genome sequenced strains obtained from the University of California at San Diego *Drosophila* stock center (now The National *Drosophila* Species Stock Center at Cornell University). The EB-GAL4 *D. melanogaster* line drives GFP in the entire EB and is a gift from Dr. Phillip Daborn (University of Melbourne). The UAS-RNAi constructs were obtained from the Bloomington Drosophila Stock Center. All flies were maintained at room temperature on standard *Drosophila* food (Bloomington formulation, Genesee Scientific). *D. melanogaster* GAL4/UAS-RNAi experiments were performed at 25 °C.

### *In situ* hybridization

Ejaculatory bulbs (EBs) from three-day old male adult flies were dissected in PBS. *In situ* hybridization was performed with RNA probes as described previously (40). Probes for *bond in situ* hybridization were synthesized from cDNA using species-specific primers (**Table S3**). The *D. melanogaster* probe was also used for *in situ* hybridization to *D. yakuba* EBs because the sequence is highly similar.

### RNA sequencing and analysis

RNA from the EBs of approximately 200 eight-day old Canton-S *D. melanogaster* males was extracted using TRIzol Reagent (Ambion, Austin, TX, USA) according to manufacturer’s instructions. Indexed RNA-Seq libraries were prepared from ~1 ug of total RNA using the TruSeq RNA Library Prep Kit v2 (Illumina) according to manufacturer’s protocol. RNA quality and concentration were measured on an Agilent 2100 Bioanalyzer (Thermo Scientific). Paired end Sequencing was performed on an NGS Illumina Hiseq 2000 with a 20 M read depth (75bp X 2; AITBiotech; Singapore). FastQ files were aligned to the Dmr6.05 *Drosophila melanogaster* reference genome (2012, r5.48) using TopHat v2.0.9 (41).

### RNAi screen

Based on the expression level of the transcription factors (TFs) in EB and predicting TF binding sites of the *bc3* fragment, 100 candidate TFs were used for the RNAi screen. Males from each *UAS::RNAi* carrying line were crossed with virgin females of the *bc3::GFP; EB-GAL4*; + fly line (**Fig. S8**). The EBs of three-day old males from the resulting crosses were dissected and imaged for GFP expression.

### Imaging

All *in situ* hybridization and GFP images were captured using the Nikon SMZ18 dissecting stereo microscope system. For GFP images, EBs were dissected from three-day old males in 1× PBS and mounted on slides with glycerol mountant [80% (vol/vol in water) glycerol, 0.1 M Tris (pH 8.0)].

## Author Contributions

J.P. and H.Ch. designed research; J.P., Z.W., H. Co, J.S.R.C., J.J., J.Y.Y., and H.Ch. performed research; J.P., J.Y.Y and H.Ch. analyzed data; and J.P. and H.Ch. wrote the paper with input from other authors.

## Acknowledgments

We thank Caitlin Peffers, Mei Luo, Ian Paulsen, and Cole Richards for technical assistance and the Bloomington Drosophila Stock Center for fly stocks and reagents. We acknowledge critical comments to the manuscript by Dr. David Arnosti (Michigan State University) and Dr. Sean B. Carroll (University of Maryland, College Park). This work is supported by USDA NIFA via Michigan State University AgBioresearch (Umbrella project MICL02522 to HC).

---

**Fig. S1.**
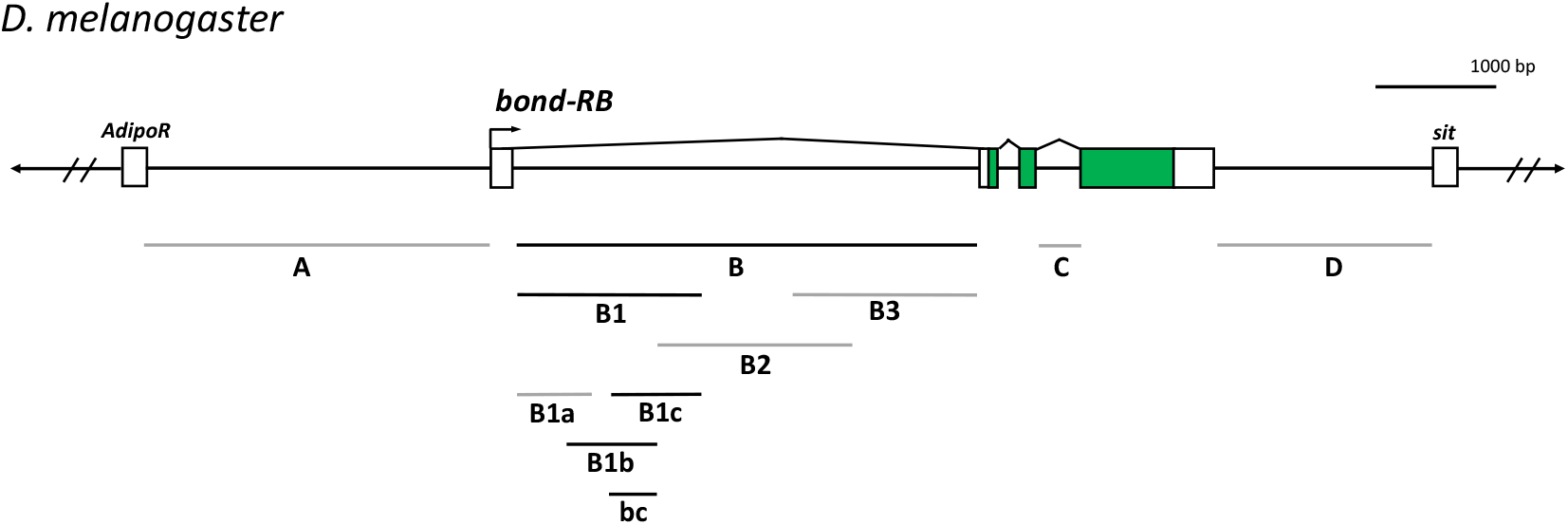
The large intron of *bond* contains sequences that can drive GFP expression in the EB semicircular wall epithelium (swe) in *D. melanogaster*. Overlapping fragments from the non-coding region around the *D. melanogaster* bond locus were screened for *cis*-regulatory sequences that were able drive GFP reporter protein expression in the EB swe. Black lines indicate fragments able to drive GFP expression in the EB swe. Grey lines indicate fragments not able to drive GFP in the EB swe.

**Fig. S2.**
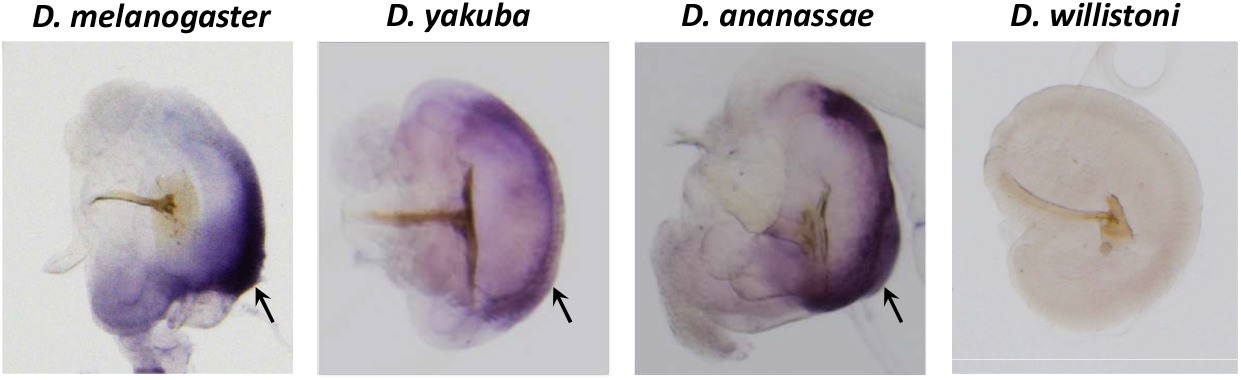
*In situ* hybridization of *bond* in the ejaculatory bulb (EB) of four *Drosophila* species. *In situ* hybridization of species-specific anti-sense *bond* probes showed that *bond* is expressed in the semicircular wall epithelium (black arrows) *of D. melanogaster, D. yakuba*, and *D. ananassae* but not *D. willistoni*.

**Fig. S3.**
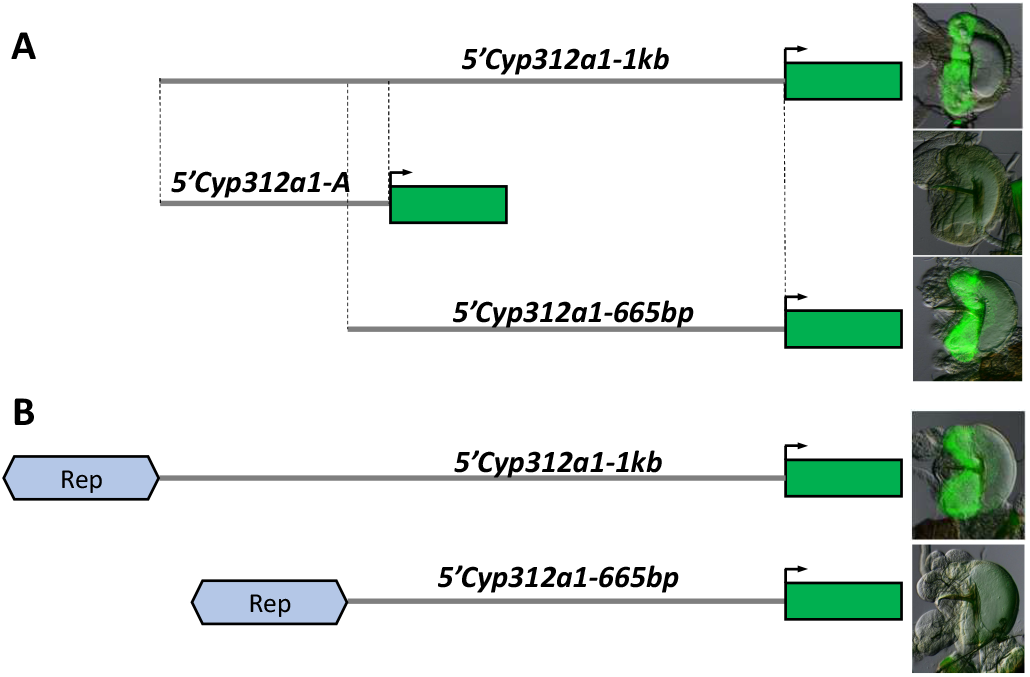
The *bond EB* repressor region (Rep) is modular and can repress gene expression of another EB enhancer in a distance dependent manner. (A) The 5’ region (1kb and 665bp) of the *Cyp312a1* gene contains enhancer sequences that drive GFP expression in the horn wall epithelium and the handle base. (B) The bond EB repressor region (Rep) can repress EB expression of the *5’Cyp312a1-665bp* fragment but not the *5’Cyp312a1-1kb* fragment suggesting that this repressor sequence may be distance dependent.

**Fig. S4.**
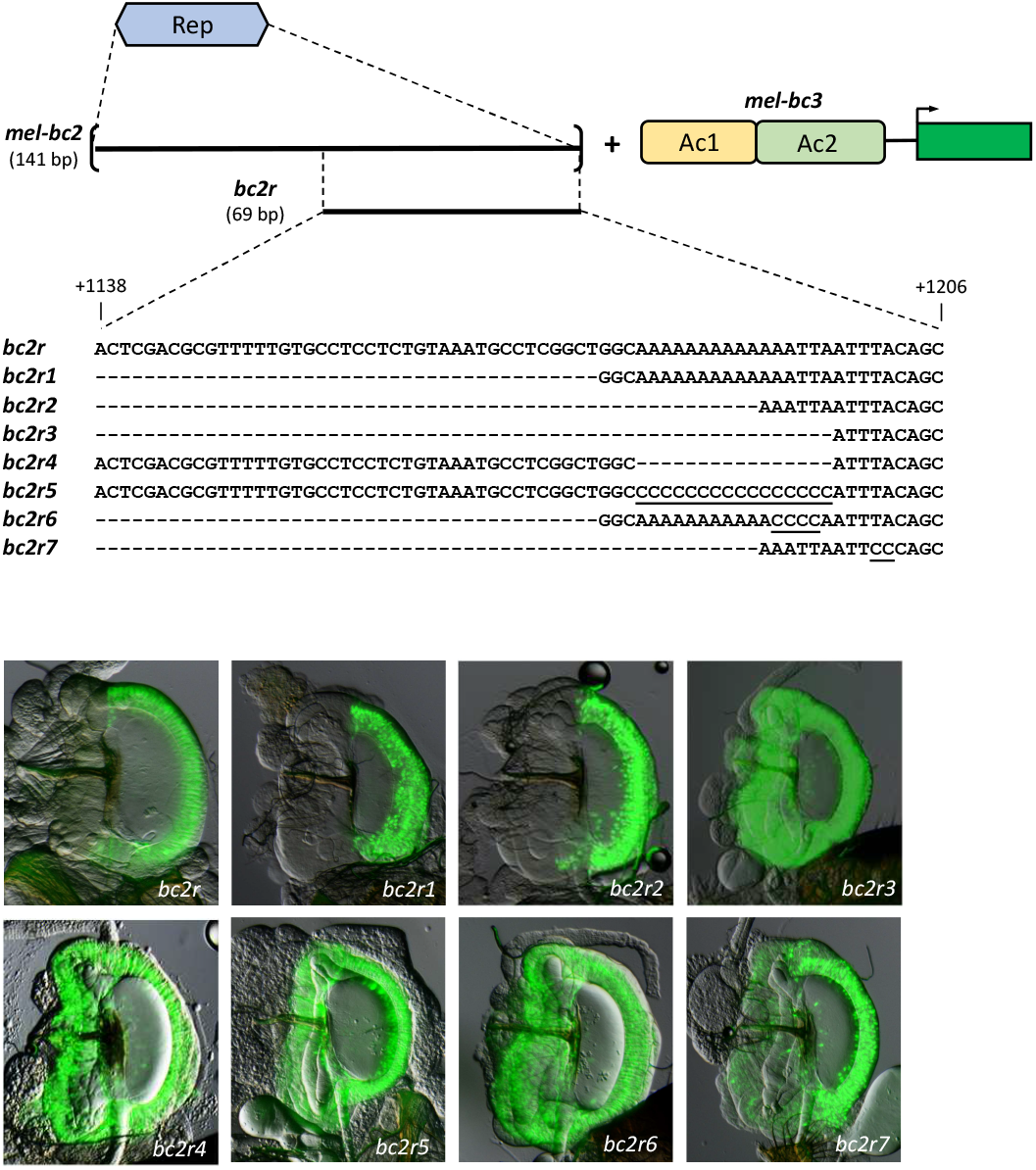
The Rep repressor can be narrowed down to a 11bp sequence in *D. melanogaster*. A series of deletion and mutation constructs are made for the 141 bp Rep region of *D. melanogaster*. The mutated nucleotides are underscored. The EB pictures show the expression of GFP driven by the different constructs. The deletion constructs bc2r, bc2r1 and bc2r2 drive same reporter expression in which GFP expression in horn wall epithelium (hwe) and handle base (hb) is repressed. The deletion constructs bc2r3 and bc2r4 drive the expression in the whole EB, fail to repress the expression in hwe and hb. bc5, bc6 and bc7 with the mutated nucleotides in different position do not repress the expression in hwe and hb.

**Fig. S5.**
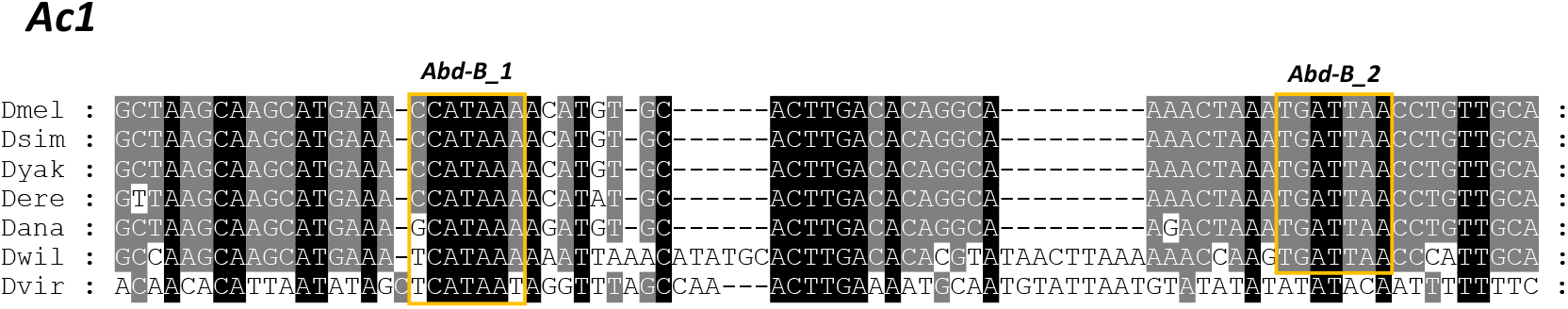
Alignment of the Ac1 region of seven *Drosophila* species. Putative *Abd-B* binding sites are indicated by the yellow boxes. The first site, *Abd-B_1*, appears to be present in all species, but the second site, *Abd-B_2*, is not present in *D. virilis*.

**Fig. S6.**
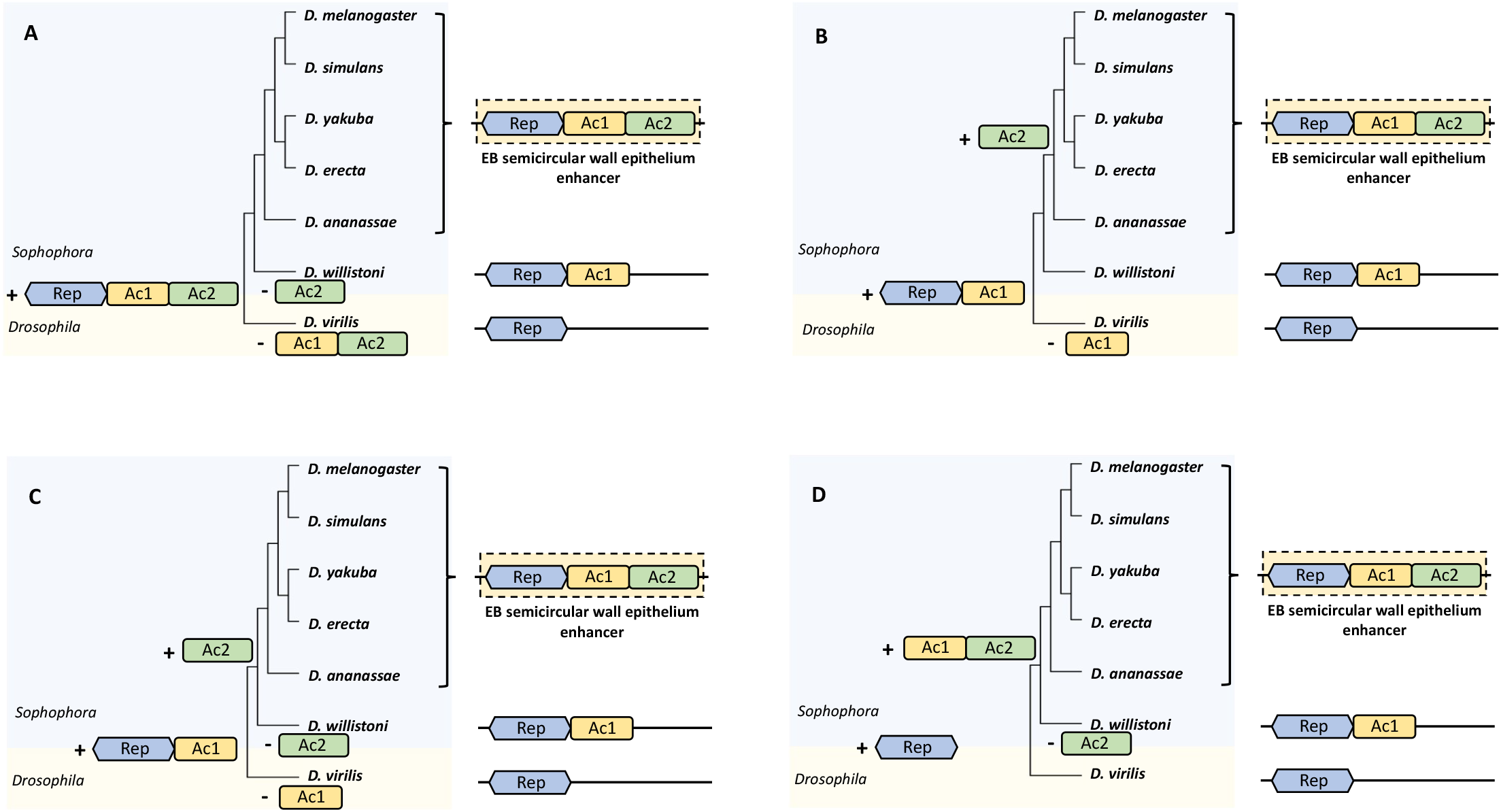
Other putative models for the evolution of EB semicircular wall epithelium enhancer. (A) This model suggest that the EB swe enhancer is ancestral to all these species and was lost in *D. willistoni* and *D. virilis* independently. However, this is unlikely as this model invoked parallel losses of Ac2 in these two species and another loss of Ac1 in *D. virilis*. (B-D) These models showed different models regarding the evolution of Rep, Ac1 and Ac2. In these three different models, the repressor precedes the gain of different activator sequences. However, these models invoked multiple gains and losses of the Ac1 and Ac2, which is less likely than the stepwise gain of these activator sequences **(Fig. 5)**.

**Fig. S7.**
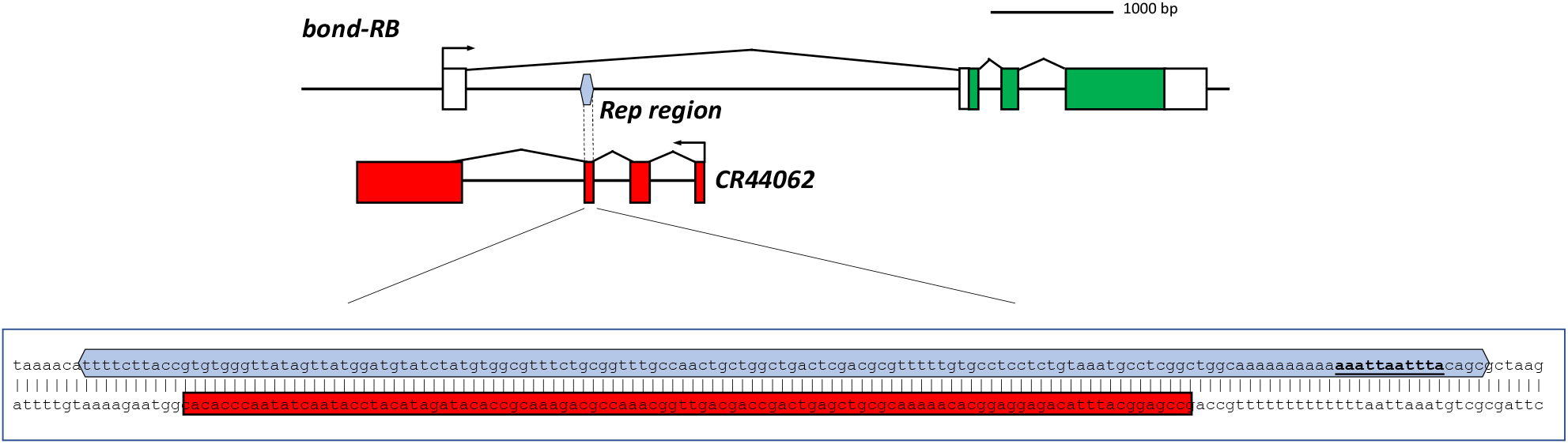
Schematic showing positions of Rep region (blue) of EB swe enhancer and the antisense non-coding RNA CR44062 relative to the *bond* locus. The Rep region overlaps with the exon of *CR44062*. The 11bp sequence is underlined and is just outside the exon of *CR44062*.

**Fig. S8.**
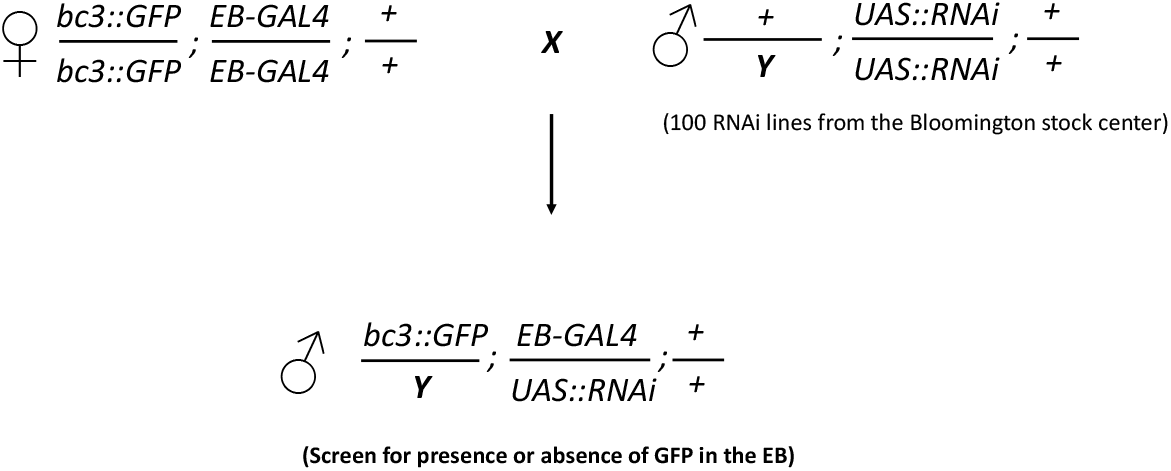
Schematic for the EB-specific RNAi screen.

## Notes

### Competing Interest Statement

The authors have declared no competing interest.

